# A systematic mutational analysis identifies a 5-residue proline tag that enhances the *in vivo* immunogenicity of a non-immunogenic model protein 240 folds

**DOI:** 10.1101/2020.04.30.070615

**Authors:** Nafsoon Rahman, Mohammad Monirul Islam, Md Golam Kibria, Satoru Unzai, Yutaka Kuroda

## Abstract

Small proteins are generally non-immunogenic, which can be a major hurdle in developing protein and peptide vaccines or producing antibodies for biopharmaceutical usage. For improving a protein’s immunogenicity, we previously proposed to use short Solubility Controlling Peptide (SCP) tags that oligomerize proteins into soluble aggregates. Here, we systematically analyzed the effect of SCP-tags that do not induce oligomerization on the immunogenicity of a small, non-immunogenic, model protein, Bovine Pancreatic Trypsin Inhibitor (BPTI-19A; 6 kDa). We assessed the effect of the following ten SCP-tags: Six tags made of five consecutive Arg, Lys, His, Asp, Asn, Pro; one made of seven Pro; two tags made of consecutive Arg-lle and Asn-Ile, all attached at the C-terminus of BPTI-19A; and a 5-proline tag attached at the N-terminus. Circular dichroism, fluorescence, dynamic light scattering measurements, and analytical ultra-centrifugation indicated that the addition of the SCP-tags did not change the secondary structure content nor the tertiary structures of the protein nor its monomeric state. On the other hand, the C-terminus 5-proline (C5P) tag unexpectedly increased the immunogenicity (IgG level) of BPTI-19A by up to 240 fold as assessed by ELISA. Additionally, the 5-arginine tag (C5R) increased the titer by up to 73 fold. The titer increase lasted for several weeks, and the effect was cumulative to that of the Freund’s adjuvant, which is commonly used to boost a protein’s immunogenicity. Altogether, SCP-tags that do not oligomerize proteins substantially increased the immunogenicity of a non-immunogenic protein, suggesting that the 5-proline and the 5-arginine SCP-tags may provide a novel tool for facilitating the production of antibodies or improving the effectiveness of protein-based vaccines.

## 1. Introduction

The poor immunogenicity of small low molecular weight proteins [1] represents a significant challenge in producing antibodies for biopharmaceutical usage as well as for developing protein-based vaccines. Adjuvants are commonly used for addressing this issue, but in biomedical practice they often encounter a patients’ reluctance to use such additives [2,3]. Besides adjuvants, virus-like particles (VLPs) have recently earned much attention in increasing the immunogenicity of proteins and a malaria vaccine candidate is being reported [4]. The fusion protein strategy, where the target protein is fused to a different protein, is another popular way for increasing immunogenicity. For example, hydrophobic lipoproteins [5], cytokines [6] or immunoglobulin domains [7] have been used to enhance the immunogenicity of target proteins. Other methods such as nano-particles [8], carrier proteins [9] and bio-conjugates [10] are also known for increasing a protein’s immunogenicity. Nevertheless, none of the above methods is perfect, and the biopharmaceutical industry is continuously in search of a novel, simple, and widely applicable method for increasing the immunogenicity of proteins.

In previous reports, we showed that short Solubility Controlling Peptide tags (SCP-tags) [11,12,13] attached at the C-terminus of a host protein could be used to induce the formation of sub-visible amorphous aggregates and thereby increase the immunogenicity of non-immunogenic proteins [14,15]. In particular, a hydrophobic isoleucine tag attached to the C-terminus of two model proteins, BPTI-19A (58 residues; MW: 5.98 kDa) and DEN3ED3 (103 residues; MW 11.46 kDa) increased their immunogenicity by oligomerizing the proteins into small nanometer-scale aggregates, and the immune response was long-lasting with a T-cell-dependent activation of the B-cells [16,17]. A major advantage of the SCP-tags is their ability to produce sub-visible aggregates in a highly reproducible and reliable way [18,19], which in turn provides good control of their effect on immunogenicity [15].

Here, we ask whether short peptide tags that increase protein solubility [20,21] and thus prevent aggregation can nevertheless increase a protein’s immunogenicity. We thus performed a systematic search for SCP-tags that would increase the immunogenicity of a non-immunogenic model protein, BPTI-19A, without oligomerizing it. To this end, we confirmed that all of the BPTI variants were monomeric and retained a native conformation, as well as biophysical properties. We then measured their immunogenicity in mice and found that C5P and C5R tags significantly enhanced the anti-BPTI-19A IgG titer, which lasted for several weeks. Thus, in addition to the previously reported C5I tag, which increased immunogenicity through protein oligomerization, C5P and C5R tags are expected to provide a tool for the effective production of biopharmaceutical antibodies and possibly for developing protein-based vaccines [14].

## 2. Materials and methods

### 2.1. Protein expression and purification

The SCP-tagged BPTI variants were constructed by introducing the SCP-tags to either the N or C termini of BPTI-19A, a simplified BPTI variant [22] containing 19 alanines out of its 58 residues [23] **(Figure 1)**. BPTI variants were named according to the number and type of amino acids added to the N or C-terminus of BPTI-19A. The DNA sequences encoding the SCP-tags were added by QuikChange site-directed mutagenesis (Stratagene, USA) using pMMHa BPTI-19A vector as a template [23], and the plasmid sequences were confirmed by DNA sequencing. The tagged variants were expressed in *Escherichia coli* BL21 (DE3) pLysS cell lines and purified according to our previously reported protocol [18].

**Figure 1.**
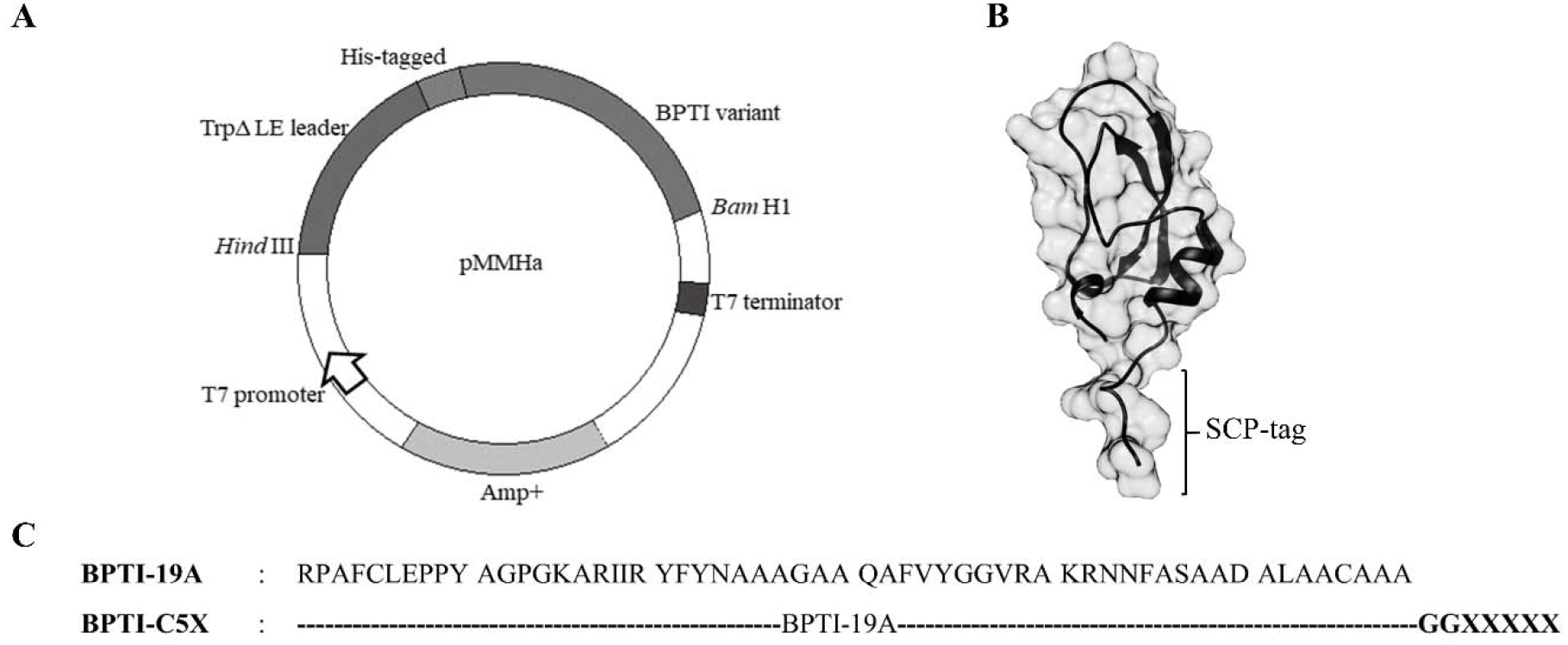
Construction and design of the tagged BPTI. A) BPTI was expressed using the pMMHa vector with a His-Trp leader. B) The tagged variants were designed by attaching the respective tags at the C-terminus, where two glycine residues served as a spacer between the target protein and the tags. The structure of SCP-tagged BPTI was generated using MODELLER (PDB ID: 3AUB) [39]. C) The sequence of BPTI-19A and the tagged variants. X stands for the one-letter amino acid code and is either R, K, N, H, D, P or I.

### 2.2. Dynamic light scattering (DLS) and static light scattering (SLS)

The hydrodynamic radii of the BPTI-19A variants were measured at 25° and 37°C by DLS on a Zetasizer Nano-S (Malvern, UK) [24]. The protein samples were prepared in PBS, pH 7.4 (filtered using 0.22 μm Minisart filter) at 0.3 mg/mL, left still for 20 minutes at 25°C and centrifuged (20,000xg for 20 minutes at 25°C). The supernatant was used for DLS and subsequent measurements as well as for immunization. Three independent readings were recorded for measuring the average hydrodynamic radius (*R*_h_) from the number distributions using the Stokes-Einstein equation [25].

The presence of sub-visible aggregates was monitored by static light scattering (SLS) at a wavelength of 600 nm using an FP-8500 spectrofluorometer (JASCO, Japan). A quartz cuvette with a 3 mm optical path length was used. Each measurement was repeated three times, and the values were averaged.

### 2.3. Analytical ultra-centrifugation (AUC)

Sedimentation velocity experiments were carried out using an Optima XL-A analytical ultracentrifuge (Beckman-Coulter, USA) with a four-hole An60Ti rotor at 33°C. AUC samples were prepared in the same way as the DLS/inoculation samples followed by overnight dialysis against PBS. PBS-density, viscosity, and BPTIs’ partial specific volumes were calculated using the SEDNTERP software [26]. Data analysis was performed using the continuous distribution *c*(s) analysis module in the SEDFIT program [27]. As a positive control, we used BPTI-C5I, which was previously reported to produce sub-visible oligomers [15,18].

### 2.4. Tyrosine fluorescence and Circular Dichroism (CD) spectroscopy

The secondary structure contents were characterized by CD spectroscopy using a JASCO J820 CD spectropolarimeter (JASCO, Japan) and spectra were measured in the wavelength range of 200-260 nm using a 1mm optical path length quartz cuvette. Tyr-fluorescence spectra with λ_ex_ 275 nm were measured on a JASCO FP-8500 spectrofluorometer using a quartz cuvette with a 3 mm optical path length. Both measurements were conducted at a protein concentration of 0.3 mg/mL in PBS, pH 7.4 at 25° and 37°C.

### 2.5. 8-anilino-1-naphthalene-sulfonic acid (ANS) fluorescence and Thioflavin T (ThT) assay

The native-like conformations of the BPTI-variants were further assessed by 8-anilino-1-naphthalene-sulfonic acid (ANS) and Thioflavin T (ThT) fluorescence measurements under conditions identical to that of the immunization experiments. Both measurements were carried out using the JASCO FP-8500 spectrofluorometer (JASCO, Tokyo, Japan) with a quartz cuvette of 3 mm optical path length at 25° and 37°C. ANS/ThT dye was added to the protein solution and incubated at 25°C for 5 minutes in the dark before measurement. The ANS-and ThT-fluorescence spectra were measured with an excitation wavelength (λ_ex_) set to respectively, 380 nm [28], and 440 nm [29]. As a positive control for amyloid formation, the hen egg-white lysozyme was used in the ThT measurement [30,31].

### 2.6. Immunization experiments

Four-week-old female mice (Jcl:ICR, CLEA, Japan) were used for the immunization experiments. For injections with adjuvant, the first dose was given subcutaneously with Freund’s complete adjuvant (WAKO, Japan), and doses 2-4 with Freund’s incomplete adjuvant, were provided intraperitoneally at weekly intervals. The quantity of injected protein was 30 μg and formulated in 100 μL of PBS or mixed with a 1:1 ratio of adjuvant when the latter was used. Control mice were injected with PBS both in the presence and absence of adjuvant. Three days after inoculation, ~20 μL of tail-bleed samples were collected and used for the weekly measurement of IgGs by ELISA. After the final dose the mice were sacrificed, heart-bleed samples were collected and stored at −30°C with 1:4 dilution in PBS supplemented with 20% glycerol.

### 2.7. ELISA

ELISA was performed as previously reported [14]. In short, anti-BPTI IgG levels were evaluated using untagged BPTI-19A (2.5 μg/mL in PBS) as coating antigen on the 96-well microtiter plates (TPP microtiter plates, Japan). Anti-BPTI sera were applied into the PBS-washed wells at an initial dilution of 1:50 and 1:300 for the tail-bleed and heart-bleed samples, respectively. Plates were then incubated at 37°C for 2 hours.

After washing the plates three times with PBS-0.05% Tween-20, each well received a 100 μL of anti-mouse IgG HRP conjugate (Thermo Fisher Scientific, USA) at 1:3000 dilution in 0.1% BSA-PBS-Tween-20 and incubated at 37°C for 1.5 hours. As a substrate, O-phenyl Di-amine (OPD) was added. The color intensities were measured at 492 nm using a microplate reader (SH-9000 Lab, Hitachi High-Tech Science, Japan) immediately after stopping the reaction with 1 N sulfuric acid (50 μL/well).

## 3. Results and Discussion

### 3.1. Biophysical characterization of the tagged BPTIs

The sizes of the tagged BPTI variants were systematically examined by dynamic light scattering (DLS) and static light scattering (SLS). DLS results showed that all tagged BPTIs were monomeric at both temperatures (**Figure 2A**, Supplementary Table S1 and Figure S1A), which was confirmed by the low scattering intensities as measured by SLS (**Figure 2C** and Supplementary Figure S1B). Analytical ultracentrifugation (AUC) also confirmed the monomeric status of all of the tagged BPTIs (**Figure 2D**, **Table 1**). Thus, all of the data indicated that the SCP-tagged BPTIs remained monodispersed and monomeric in solution alike the untagged BPTI-19A (**Figure 2A-2D**). As control of an aggregated mutant, we used the C5I-tagged BPTI, which formed oligomers with a distribution of sizes (**Figure 2A-2D**), in line with our previous report [15,18]. In addition, DLS was measured before each round of injection (*i.e.* a near ‘real-time monitoring’ of *R*_h_) in order to fully ensure that the tagged samples (except BPTI-C5I) were actually monomeric at the time of injection (**Figure 2B**).

**Figure 2.**
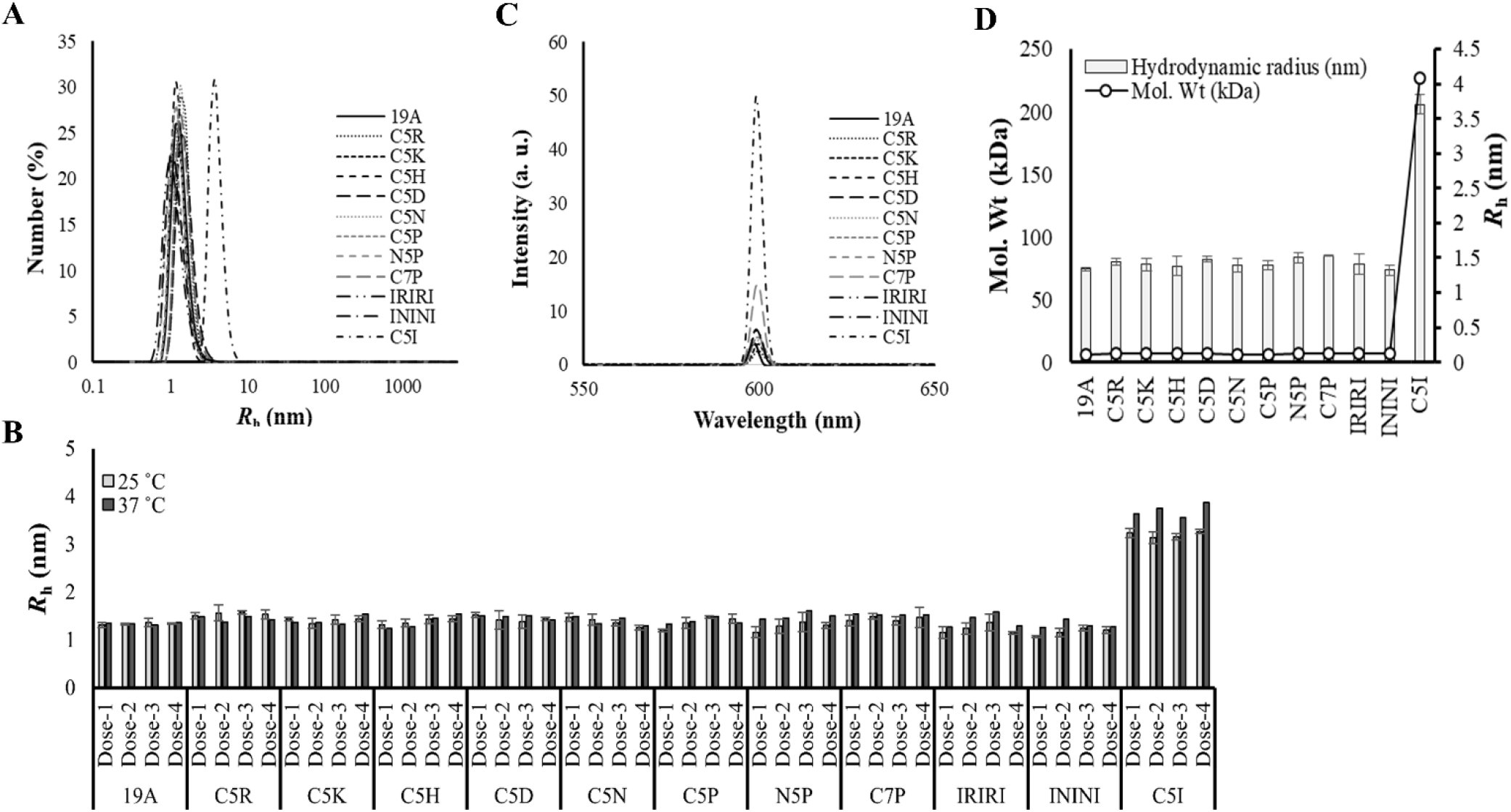
Analysis of the monomeric and oligomeric status of the tagged and untagged-19A under physiological conditions. A) Number distribution of the hydrodynamic radii (*R*_h_) measured by DLS. B) *R*_h_ measured just before immunization from dose-1 to dose-4 and computed from DLS’s number distribution. C) SLS spectra of the BPTI variants. D) Molecular weights of tagged variants measured at 33°C by AUC and their correlation with the *R*_h_. DLS/SLS data were the averages of three independent measurements. Line symbols are explained within the panels.

**Table 1:** Molecular weights of BPTI-19A variants by Analytical Ultra Centrifugation (AUC)/ Sedimentation velocity. AUC experiments were carried out with a protein concentration of 0.3 mg/mL in PBS at 33°C. The *c* (s) distribution was converted into a molar mass distribution *c* (M), from which the average molecular weights were calculated. *The molecular weight of BPTI-C5I could not be determined as it formed oligomers with multiple sizes.

| **Mutant Identities** | **Molecular weights (kDa)** | **Population (%)** |
| --- | --- | --- |
| BPTI-19A | 6.1 kDa | 90.1% |
| BPTI-C5R | 7.0 kDa | 98.1% |
| BPTI-C5K | 6.9 kDa | 96.2% |
| BPTI-C5H | 6.7 kDa | 95.2% |
| BPTI-C5D | 6.8 kDa | 97.2% |
| BPTI-C5N | 6.1 kDa | 74.7% |
| BPTI-C5P | 6.4 kDa | 97.2% |
| BPTI-N5P | 6.6 kDa | 91.7% |
| BPTI-C7P | 6.9 kDa | 90.7% |
| BPTI-IRIRI | 6.6 kDa | 95.0% |
| BPTI-ININI | 6.5 kDa | 94.7% |
| BPTI-C5I | Large* | * |

Circular dichroism (CD) indicated that the tagged BPTI variants had a secondary structure content similar to that of the untagged BPTI-19A, and Tyr-fluorescence indicated that they retained a native-like tertiary structure (**Figure 3A-3B** and Supplementary Figure S1C-1D). Furthermore, ANS-fluorescence intensity of the untagged-19A and all other variants remained very low compared to that of BPTI-C5I **(Figure 3C** and Supplementary Figure S1E), confirming their native-like, non-molten globular structural properties. Moreover, no ThT-fluorescence signal was observed indicating the absence of amyloidogenic aggregates [28,32] (Supplementary Figure S1F). Altogether, all of the measurements indicated that none of the tags (except the previously reported C5I) affected the conformation nor the monomeric state of BPTI-19A under injecting conditions.

**Figure 3.**
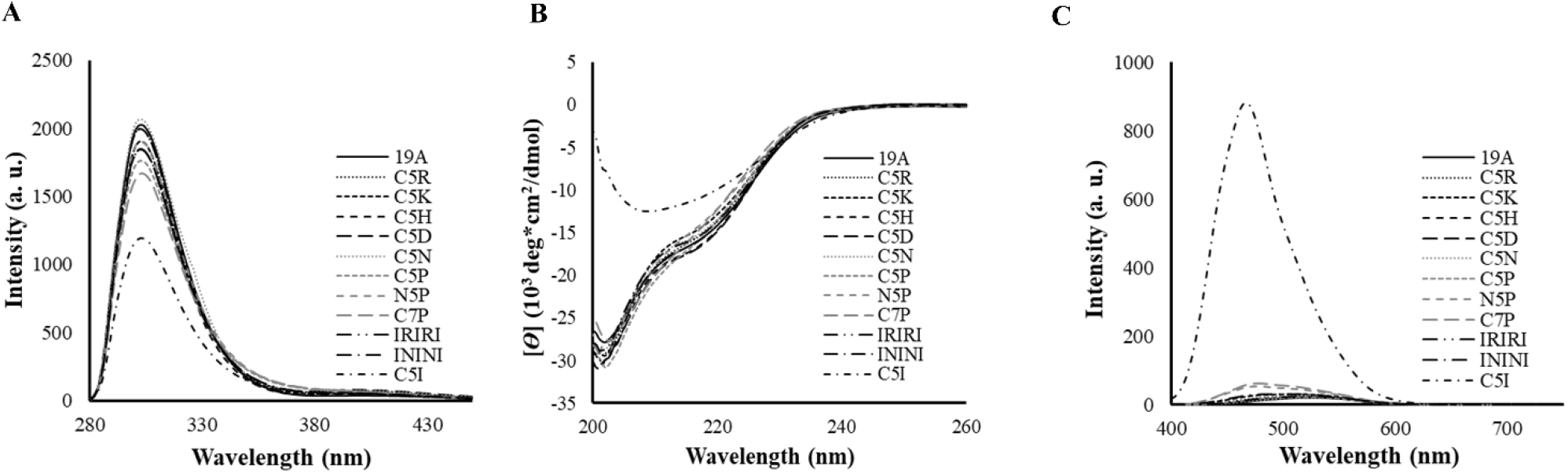
Structural characterization of the BPTI-19As under physiological conditions. A) Tyr-fluorescence, B) CD, and C) ANS-fluorescence spectra measured at a protein concentration of 0.3 mg/mL in PBS, pH 7.4. Data are the averages of three accumulations. Line symbols are explained within the panels.

### 3.2. Search for a non-aggregating immunogenicity-enhancing peptide tag

We investigated the immunogenicity of all of the ten tagged BPTI-19A in Jcl:ICR mice model both in the presence and absence of adjuvant. The untagged BPTI-19A, which remained monomer, was non-immunogenic as expected for a small 6 kDa protein (**Figure 4**), even in the presence of Freund’s adjuvant. However, the antibody titers of some of the tagged BPTI-19As became noticeable after the third injection (Supplementary Figure S2), and after the final dose (4^th^), the titer of the anti-BPTI-C5P was 240 fold higher than that of the untagged BPTI-19A (**Figure 4A**, Supplementary Figure S3A). This increase was even higher than the 182 fold increase observed in our control experiment, where the oligomerizing C5I tag was attached to BPTI-19A (**Figure 4A** and **Table 2**). To date, the C5R tag, which also kept BPTI-19A monomeric at the time of injection, increased the immunogenicity of BPTI-19A by 73 fold. Interestingly, N5P barely increased BPTI-19A’s immunogenicity, and C7P increased the immune response by 149 fold, which was a substantial increase but less than we expected from the increase resulting from the C5P tag (**Figure 4A** and **Table 2**).

**Figure 4.**
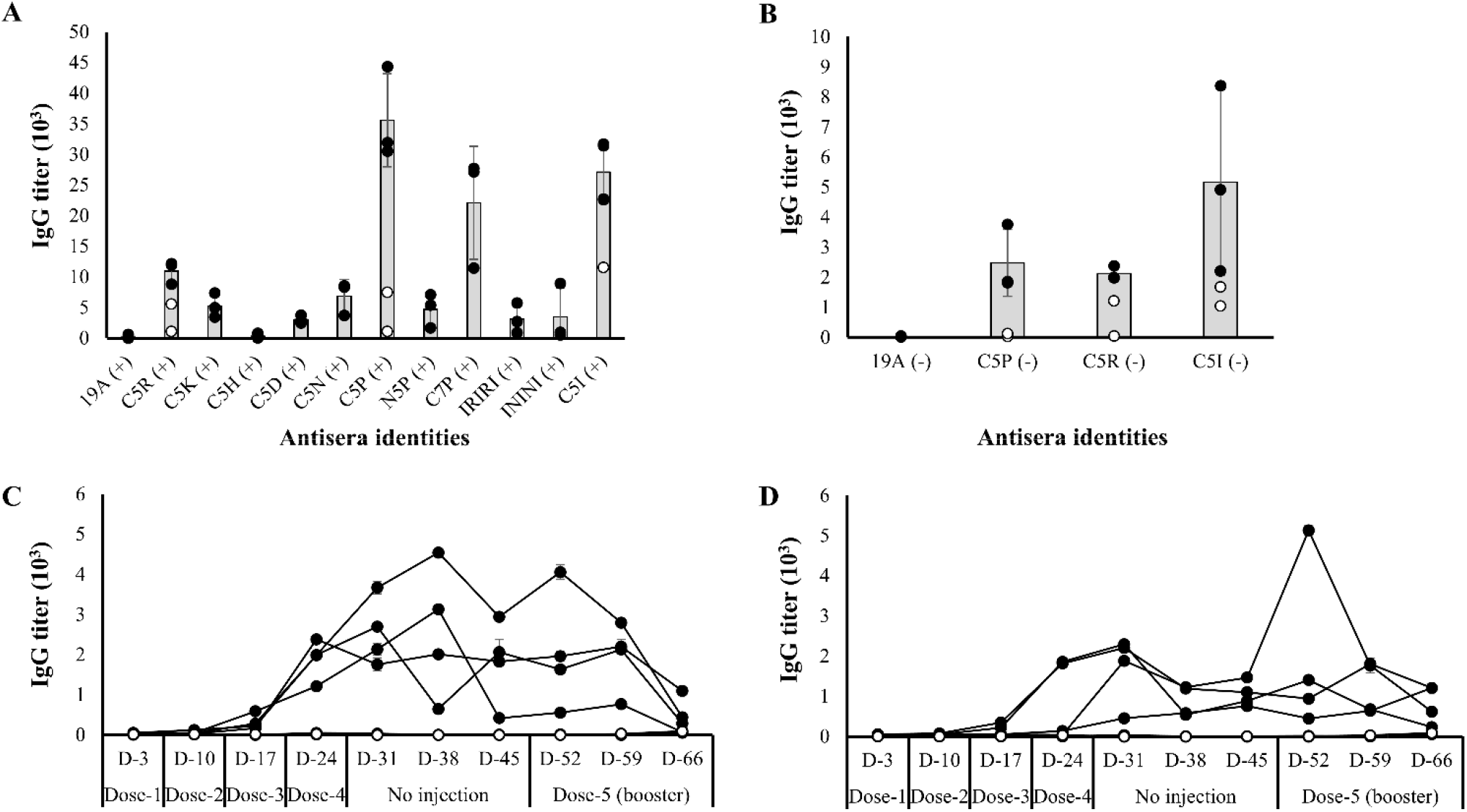
Antibody responses against the tagged and the untagged BPTI-19A. Immunization in the presence and absence of adjuvant are indicated, respectively, by ‘+’ and ‘-’. A) IgG titers after the final dose (4^th^ dose) were measured using the tail-bleed sera of mice in the presence of adjuvant. Outliers (open circles) were removed when computing the average IgG titer (grey bars), which was calculated using data from the high responsive mice (closed circles) [19A (+), n=4, C5P/R/I (+), n=5, C5K/D/N/H/IRIRI/ININI (+), n=3, N5P/C7P (+), n=3; ‘n’ indicates the number of mice]. B) Antibody titers of BPTI-19A, BPTI-C5P, BPTI-C5R and BPTI-C5I in the absence of adjuvants after the final (4^th^ dose) using the tail-bleed sera. The average IgG titer (grey bars) was calculated using data from high responsive mice (closed circles) and the outliers were excluded (open circle) [19A (−), n=3; C5P/R/I (−), n=5]. Long term IgG titers against C) untagged-19A (open circle), BPTI-C5R (closed circles) and D) BPTI-C5P (closed circles) in the absence of adjuvant [19A (−), n=3 and C5P/R (−), n=4]. ‘D’ indicates the day at which tail bleeding was performed counting from the first inoculation. A booster dose was given on D-45 and the IgG titer on D-52 indicates the immune response a week after the booster dose was administrated. Line symbols are explained within the panels.

**Table 2:**
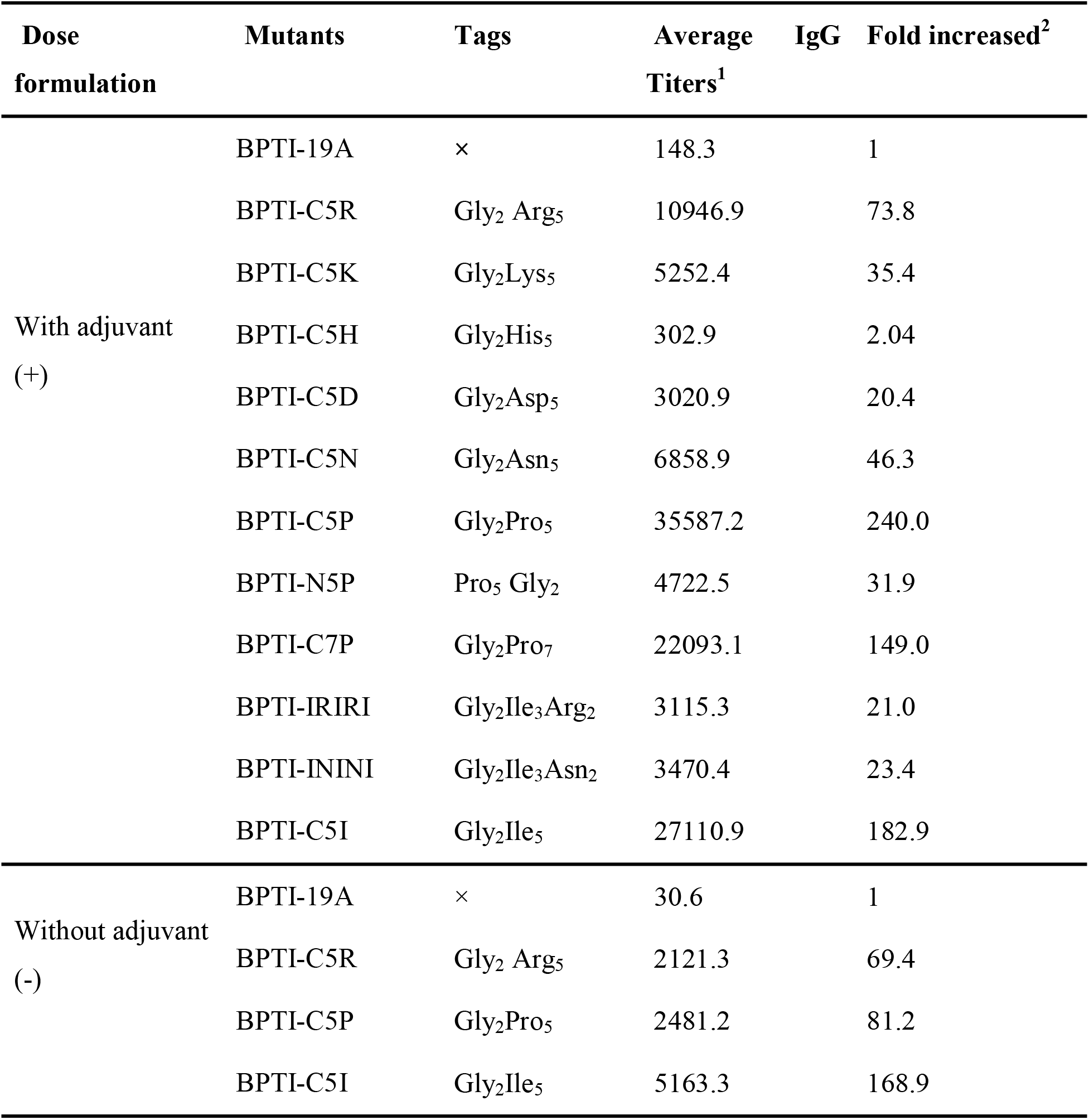
Average IgG titer values for tagged BPTI variants with and without adjuvant. ^1^The titers were calculated using a power fitting model, and the values were averaged using the number of highest responsive mice in respective groups [with adjuvant group: 19A (+), n=4, C5P/R/I (+), n=5, C5K/D/N/H/IRIRI/ININI (+), n=3, N5P/C7P (+), n=3; without adjuvant group: 19A (−), n=3; C5P/R/I (−), n=5; where ‘n’ indicates the number of mice]. ^2^The fold-increase of the titers in the presence and absence of adjuvant was calculated with respect to the titer of BPTI-19A (+) and BPTI-19A (−), respectively. ‘×’ indicates the absence of tags.

Immunization experiments in the absence of adjuvant were carried out for tags that showed the highest immune response. Namely, the C5P and C5R-tagged BPTI-19As, the untagged BPTI-19A and the oligomerizing C5I-tagged BPTI-19A were used. Noteworthy and in line with our previous observation, the C5I tag increased the antibody titer of BPTI-19A by 168 fold, reflecting the importance of the nanometer size aggregates in increasing a protein’s immunogenicity (**Figure 4B** and **Table 2**). C5P and C5R tags significantly increased the immunogenicity of the BPTI-19A by 81 and 69 fold, respectively. Importantly, all of the above ELISA figures remained essentially unchanged whenever the plates were coated with the tagged-BPTIs (own-tags) or the untagged BPTI-19A (Supplementary Figure S3C) indicating that the IgGs were raised against BPTI-19A itself and not the tags. Furthermore, the effect of the tags was cumulative to that of the Freund’s adjuvant, indicating its strong potential for acting as a target-specific adjuvant (Supplementary Figure S3B).

Finally, we assessed the maintenance of the immune responses generated by the untagged-19A, BPTI-C5P, and BPTI-C5R, and monitored the IgG level for several weeks after the last injection. Mice were weekly injected four times with the corresponding variant in the absence of adjuvant and kept untreated for three weeks, during which the IgG titer was measured weekly through tail-bleeding. Anti-BPTI-19A IgG levels for the C5P and C5R tagged BPTI-19A sustained during the three weeks (**Figure 4C-4D**), and a booster dose (5^th^ dose) of the respective variants injected twenty-one days after the 4^th^ dose increased the IgG titers by 334 and 346 fold, respectively (Supplementary Table S2). On the other hand, the IgG titer of mice immunized with the untagged BPTI-19A remained nil throughout the entire immunization period (**Figure 4C-4D**). Thus, in the absence of adjuvant, the immunogenicity increase generated by the non-aggregating BPTI-C5P and BPTI-C5R were not as outstanding as that of the BPTI-C5I [15] but nevertheless significant, and the higher titers were maintained for several weeks (**Figure 4B-4D**).

### 3.3. Short peptide tag: a biotechnological tool for increasing immunogenicity

This study was initiated in order to complement our previous ones, where we demonstrated that protein’s immunogenicity could be boosted by attaching an SCP-tag that oligomerizes the protein. Here, the immunogenicity improvement induced by the proline and the arginine tags probably involve mechanisms distinct from that generated by the C5I tag, which acts through the oligomerization of the antigens [16,17]. However, from an application viewpoint, the critical finding is the sharp increase of BPTIs’ immune response generated by the C5P tag. On the other hand, 5 prolines attached at the N-terminus (N5P) had virtually no effect on the immunogenicity of BPTI-19A, and the C-terminus 7-proline (C7P) increased the immunogenicity to a lesser extent than expected from the C5P tag.

Thus, we hypothesize that the molecular mechanisms of the C5P-induced immunogenicity enhancement are different from the previously reported high immunogenicity of proline-rich epitopes because the epitopes were located within the protein [33,34] which contrast with our observation that the immune response increased only with the 5-pro attached at the C-terminus of the protein. We also hypothesize that the 5-arg might have enhanced the entry of BPTI into the cells during the endocytosis/phagocytosis process [35,36,37], which might have increased the immunogenicity [38] of BPTI. Finally, the immunogenicity increase generated by the IRIRI and ININI were lower than C5R, C5N or C5I, indicating that the effect of the type of amino acid (contained in the tags) on immunogenicity was not additive.

## 4. Conclusions

To the best of our knowledge, this is the first systematic analysis of the effects of short peptide tags on the immunogenicity of a small and non-immunogenic protein. We found that the C5P and C5R tags could considerably increase the immune response against BPTI-19A without oligomerizing the protein. The immunogenicity improvement induced by the C5P and C5R tags probably involve mechanisms distinct from that generated by the C5I tag, which acts through the oligomerization of the antigens

## Abbreviations

BPTI: Bovine Pancreatic Trypsin Inhibitor
CD: Circular Dichroism
DLS: Dynamic Light Scattering
SLS: Static Light Scattering
AUC: Analytical Ultra Centrifugation
SCP-tags: Solubility Controlling Peptide tags
ELISA: Enzyme-Linked Immunosorbent Assay

## Acknowledgments

We thank Mr. Yuki Matsuzawa and all members of the Kuroda’s laboratory for technical help and discussion. We are grateful to Profs Tsuyoshi Tanaka and Tomoko Yoshino and for the use of the Zetasizer.

