## Supplementary Information for "A systematic mutational analysis identifies a 5-residue proline tag that enhances the *in vivo* immunogenicity of a non-immunogenic model protein 240 folds"

^1^Department of Biotechnology and Life Sciences, Graduate School of Engineering, Tokyo University of Agriculture and Technology, 2-24-16 Nakamachi, Koganei-shi, Tokyo 184-8588, Japan. ^2^Department of Biochemistry and Molecular Biology, University of Chittagong, Chittagong-4331, Bangladesh, ^3^ Department of Frontier Bioscience, Faculty of Bioscience and Applied Chemistry, Hosei University, 3-7-2 Kajino-cho, Koganei, Tokyo 184-8584, Japan.

**Supplementary Information**

**Figure S1:** Analysis and comparison of biophysical properties of C5R, C5K, C5N, C5H, C5D, C5P, N5P, C7P, IRIRI, ININI and C5I tagged BPTIs with untagged-19A. (A) DLS spectra at 25 °C of the size distribution shown as number mean. (B) SLS spectra of BPTI variants at 25°C. (C) CD spectra of all BPTI variants at 25°C. (D) Tyr-fluorescence and (E) ANS-fluorescence spectra of untagged-19A, C5R, C5K, C5N, C5H, C5D, C5P, N5I, C7P, IRIRI, ININI and C5I tagged variants at 25°C. (F) ThT-fluorescence intensity of the tagged variants measured at 37°C where lysozyme was used as a positive control. SCP-tagged variants were formulated at 0.3 mg/mL concentrations in PBS, pH 7.4 for all measurements. Values are shown as the average of three independent measurements (DLS) and three accumulations (SLS, CD and fluorescence), respectively. Line symbols are explained within the panels.

**Figure S2:** Dose-dependent titer values of all mice injected with C5R, C5K, C5N, C5H, C5D, C5P, N5P, C7P, IRIRI, ININI and C5I tagged BPTIs and untagged BPTI-19A (A) in the presence (+) and (B) absence (-) of adjuvant.

**Figure S3:** (A) Dose-dependent (with adjuvant) OD values of C5R, C5K, C5N, C5H, C5D, C5P, IRIRI and ININI tagged variants and the untagged BPTI-19A at 492 nm determined by ELISA using the 4^th^ tail-bleeding serum samples. Values are the average of duplicated samples. Line symbols are explained within the panels. (B) Comparison of antibody titers of C5P, C5R, and C5I tagged BPTIs and the untagged-19A injected with or without adjuvants. Outliers (open circles) were removed when computing the average IgG titer (grey bars), which was calculated using data from the high responsive mice (closed circles). (C) OD values of sera (using adjuvants) raised against untagged-19A, C5R and C5P tagged BPTIs against the coating antigens of untagged BPTI-19A and their respective tag, self-tags.

**Table S1: Hydrodynamic radius of BPTI-19A and its SCP-tagged variants.** DLS measurements were carried out just before immunization at 25°C followed by 37°C at 0.3 mg/mL concentrations in PBS. The values are averaged over three independent measurements, and the errors are standard deviation (SD). *R*_h_ was calculated from the number-distributions using the Stokes-Einstein equation.

**Table S2: Maintenance of IgG titers against BPTI-19A, BPTI-C5R, and BPTI-C5P in the absence of adjuvant.** ^1^ The titers were calculated using a power fitting model, and the values were averaged using the number of the mice (n) in the respective groups [19A (-), n=3 and C5R (-), n=4, and C5P (-), n=4]. ^2^Fold-increase with respect to the titer of BPTI-19A.

**Table S1:** **Hydrodynamic radius of BPTI-19A and its SCP-tagged variants**

| **Mutant Identities** | **Average hydrodynamic radius (*R*_h,_ nm) in PBS** | |
| --- | --- | --- |
|  | 25°C | 37°C |
| BPTI-19A | 1.33±0.02 | 1.34±0.02 |
| BPTI-C5R | 1.54±0.03 | 1.44±0.05 |
| BPTI-C5K | 1.41±0.05 | 1.4±0.09 |
| BPTI-C5H | 1.39±0.06 | 1.38±0.14 |
| BPTI-C5D | 1.44±0.06 | 1.48±0.04 |
| BPTI-C5N | 1.38±0.09 | 1.4±0.09 |
| BPTI-C5P | 1.37±0.09 | 1.4±0.07 |
| BPTI-N5P | 1.28±0.08 | 1.5±0.08 |
| BPTI-C7P | 1.44±0.04 | 1.53±0.09 |
| BPTI-IRIRI | 1.23±0.09 | 1.41±0.15 |
| BPTI-ININI | 1.16±0.07 | 1.32±0.07 |
| BPTI-C5I | 3.2±0.06 | 3.71±0.13 |

**Table S2: Maintenance of IgG titers against BPTI-19A, BPTI-C5R, and BPTI-C5P in the absence of adjuvant**

| **Mutants** | **Tags** | **Average titer after booster dose (D-52) ^1^** | **Fold increased^2^** |
| --- | --- | --- | --- |
| BPTI-19A | **×** | 5.92 | 1 |
| BPTI-C5R | Gly_2_ Arg_5_ | 2053.25 | 346.55 |
| BPTI-C5P | Gly_2_Pro_5_ | 1981.96 | 334.52 |

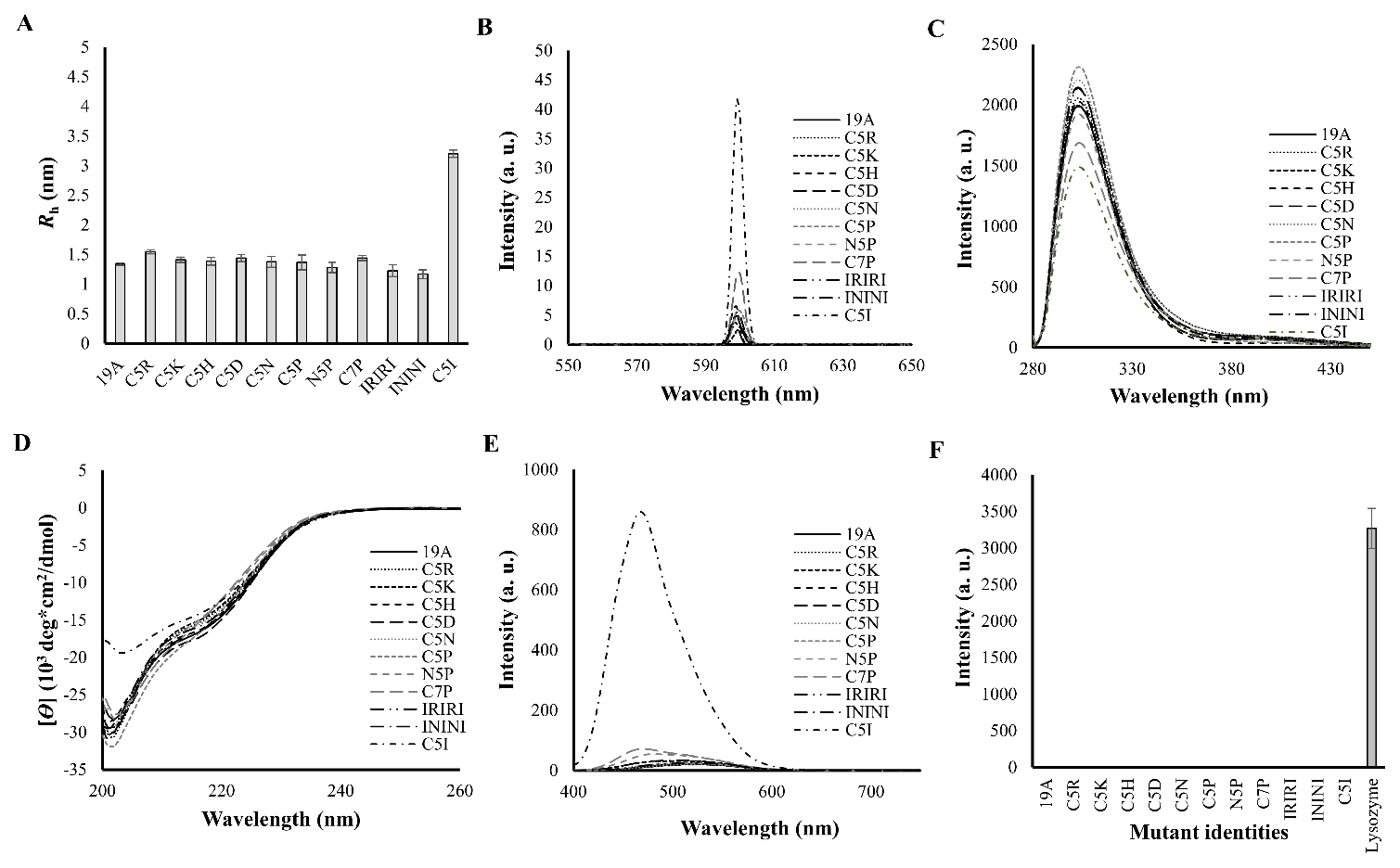
**Figure S1**

**Figure S2**

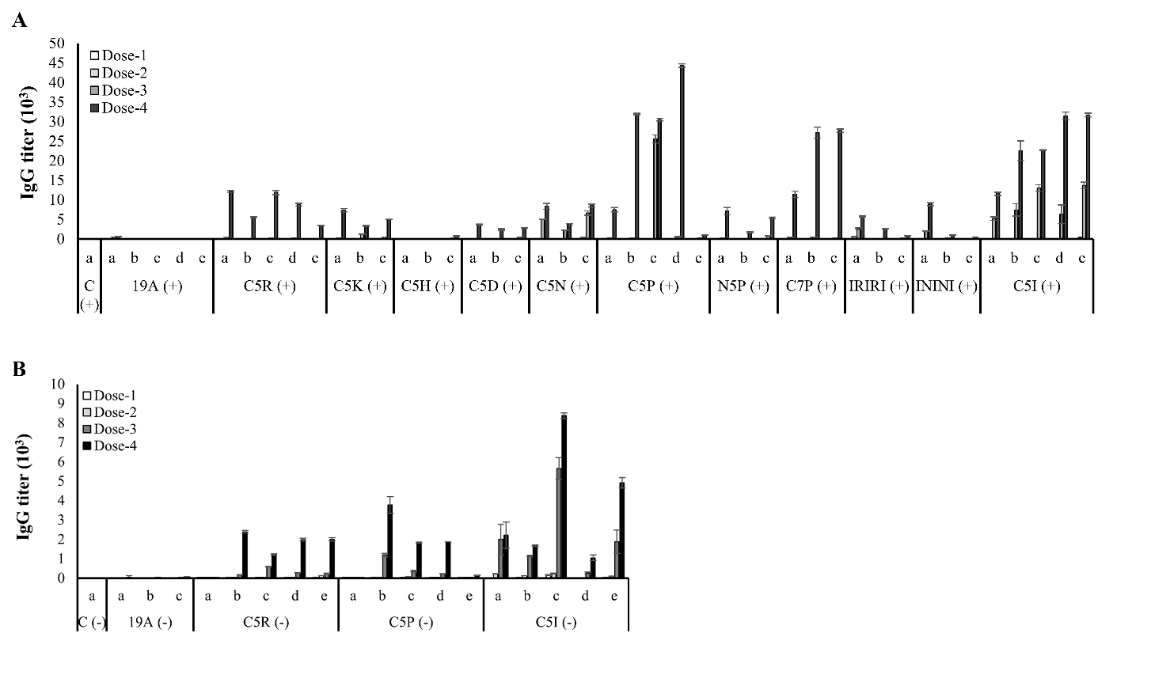

**Figure S3**

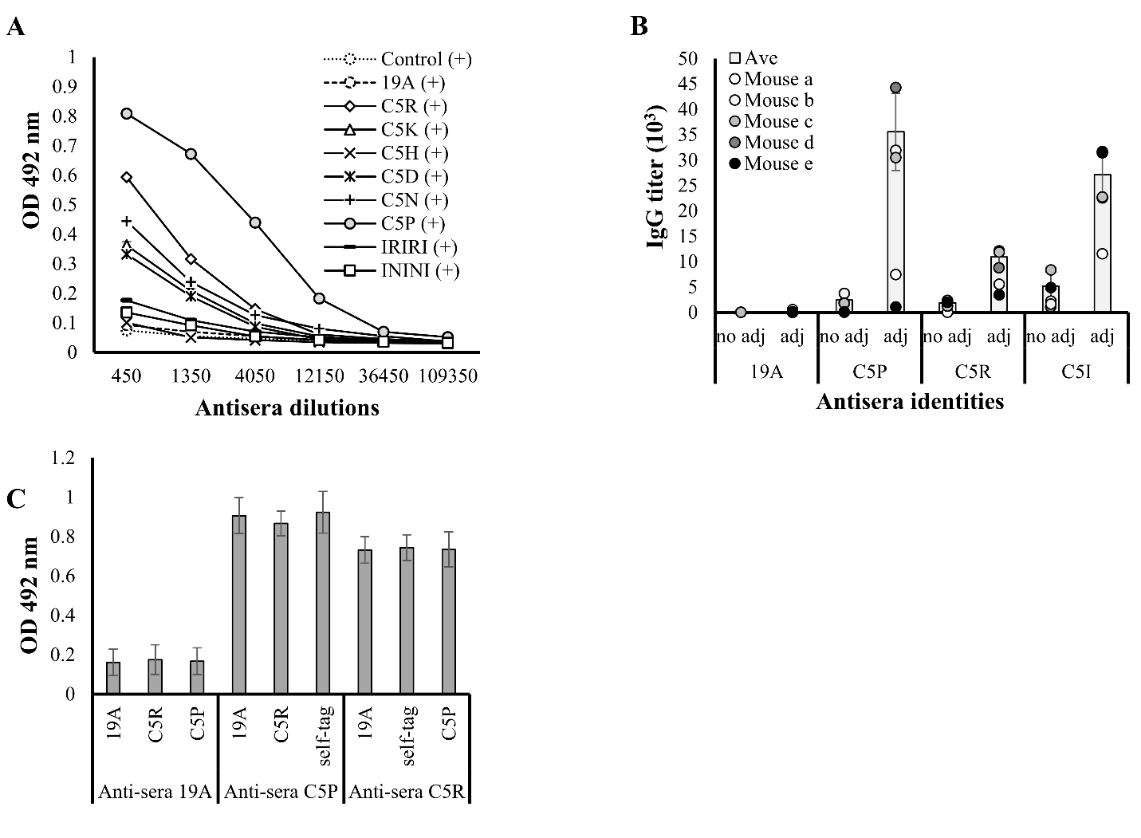
